# Ubiquitin ligase PUB41 modulates root hair development in Arabidopsis via interaction with the auxin polar transporter PIN2

**DOI:** 10.64898/2025.12.25.696480

**Authors:** Avinash Sharma, Dina Raveh, Dudy Bar-Zvi

## Abstract

Root hairs are short-lived, elongated tubular extensions of epidermal cells, that increase the surface area improving absorption of water and nutrients. Root hair development is affected by developmental, environmental, and hormonal cues, primarily Auxin. Auxin is synthesized in shoot and root apical transporters. Here, we report that mutants of the Arabidopsis *PLANT U-BOX41* (*AtPUB41*) gene encoding a ubiquitin ligase (E3) develop fewer and shorter root hairs than wild-type (WT) plants. *pub41* mutants are not impaired in sensing and responding to exogenous auxin, suggesting that the root hair phenotype may result from a change in auxin levels and/or distribution. Indeed, Auxin enhances the steady-state levels of *PUB41* transcript and protein in the roots. A catalytically inactive decoy protein lacking the U-box, PUB41(ΔU), complements the sparse root hair phenotype of *pub41* mutants to the same extent as full-length *PUB41.* Plants expressing *PUB41(ΔU*) are hyperresponsive to auxin-induced root hair development and PUB41(ΔU) accumulates in the root elongation and differentiation zones, preferentially in trichoblast cells and their root hairs. Furthermore, in these cells, PUB41-eGFP and PUB41(ΔU)-eGFP are primarily located on the anticlinal faces, as is the auxin PIN2 exporter. We show colocalization of PUB41(ΔU)-eGFP and PIN2-mCherry, and binding *in vitro* of PUB41 to the cytoplasmic hydrophilic loop of PIN2. Moreover, PUB41(ΔU) protein stabilizes PIN2 and enhances its abundance in the roots. Our results indicate that PUB41 modulates root hair development via interaction with PIN2.

**Significance Statement:** Root hairs are extensions of the root epidermal cells, which are crucial for the uptake of water and nutrients from the soil. Root hair development is highly regulated and affected by developmental, environmental, and hormonal cues. In this study, we demonstrate that the auxin-induced ubiquitin ligase, PLANT U-BOX 41 (PUB41), regulates root-hair development by interacting with the auxin polar transporter PIN2 responsible for determining auxin levels.

## Introduction

Roots anchor the aerial plant organs, absorb water and minerals and their architecture is tightly controlled by both developmental and environmental cues (reviewed by Petricka *et al*., 2012). Roots extend by cell division at the root tip meristem (Bibikova and Gilroy, 2002). The new cells elongate vertically in the elongation zone and then undergo specialization in the differentiation zone (Dolan and Davies, 2004; Verbelen *et al*., 2006). Tubular protrusions extend perpendicular to the primary root axis from trichoblastic epidermal cells, forming unicellular root hairs (Dolan and Davies, 2004). These short-lived structures increase root surface area, enhancing water and mineral uptake (Tanaka *et al*., 2014). Root hair pattern development in Arabidopsis is defined as type 3 (Cui *et al*., 2018): the epidermal cells are positioned over an inner layer of cortical cells leading to alternating columns of trichoblasts and atrichoblasts (Kim *et al*., 2006; Bibikova and Gilroy, 2002; Kim *et al*., 2006; Cui *et al*., 2018).

Auxin and ethylene are the principal positive regulators of root hair development (Vissenberg *et al*., 2020), with ethylene interacting with auxin signaling (Dubois *et al*., 2018). Auxin regulates different aspects of root development including root hair and lateral root development, cell division, expansion, differentiation, and tropic responses (Friml, 2022). Auxin is transported in a polarized manner, resulting primarily from the activity of auxin uptake and efflux via AUXIN RESISTANT1/LIKE AUXIN (AUX1/LAX) and PIN-FORMED (PIN) family transporters, respectively (reviewed by Hammes *et al*., 2022). PIN transporters act as functional dimers, each monomer consists of 10 membrane-spanning helices with cytoplasmic hydrophilic loops between helices 5 and 6 (Joshi and Napier, 2023). PIN transporters are distributed in a polar fashion in many cell types, resulting in directional auxin transport. In particular, PIN2 is localized at the basal side of young cortex cells and the apical side of root epidermal cells (Michniewicz *et al*., 2007) and is involved in root hair development and gravitropism: both functions are impaired in *pin2* mutants (Rigas *et al*., 2013). Interestingly, overexpressing *PIN2* driven by the root hair-specific *EXPA7* promoter inhibits root hair growth (Ganguly *et al*., 2010). The PIN proteins’ hydrophilic loop has residues that act as cues for posttranslational modification that regulate their polarity, activity, and trafficking (Michniewicz *et al*., 2007). Differential ubiquitylation of PIN2 acts as a signaling cue regulating its activity, stability, and protein localization, which in turn affects polar auxin distribution and the overall growth response (Abas *et al*., 2006; Leitner *et al*., 2012).

We recently reported that Arabidopsis *PUB41* (At5g62560) has a central role in seed dormancy, germination, and response to drought, and acts as a positive regulator of abscisic acid (ABA) (Sharma *et al*., 2024). *PUB41* accumulates in root tissue due to elevated promoter activity in root hairs and root meristems. PUB41 belongs to the small Plant U-Box (PUB) subfamily of ubiquitin ligases (Azevedo *et al*., 2001; Wiborg *et al*., 2008; Trujillo, 2018). with multiple Armadillo motifs that facilitate protein-protein interactions involved in target selection, alongside the U-box motif that binds the ubiquitin-conjugating enzyme. PUB-ARM E3s function as hubs of the stress response (Trenner *et al*., 2022).

Here we present evidence for the involvement of *PUB41* in root hair development. T-DNA insertion mutants of *pub41* exhibit impaired root hair development, which is rescued by the expression of either full-length PUB41 or a catalytically inactive PUB41(ΔU) protein lacking the U-box domain. *pub41* mutants are not impaired in sensing and responding to exogenous auxin, suggesting that the root hair phenotype may result from a change in auxin levels and/or distribution. Plants expressing *PUB41(ΔU*) are hyperresponsive to auxin-induced root hair development. PUB41(ΔU) accumulates in the root elongation and differentiation zones, preferentially in trichoblast cells and their root hairs. In the meristem and elongation zones, the PUB41 protein is found throughout the nucleus and cytosol. In contrast, in the upper zones of the roots, cytoplasmic PUB41-eGFP is primarily enriched at the anticlinal faces of the outer layers of the root, in a pattern resembling that of PIN auxin exporters (Michniewicz *et al*., 2007). Indeed, PUB41-eGFP and PUB41(ΔU)-eGFP colocalized with PIN2-mCherry *in vivo* and *in vitro* via interaction between PUB41(ΔU) and the cytoplasmic hydrophilic loop of PIN2. Our results indicate that the interaction between PUB41 and the auxin polar transporter PIN2 is important for root hair development.

## Results

### *pub41* mutants have fewer root hairs

Under regular growth conditions, *pub41* mutant plants look similar to WT plants (Sharma *et al*., 2024). However, on close examination, *pub41* seedlings show a significant reduction in the density and length of the root hairs (**Figure 1A-C**). In addition, in the elongation and differentiation zone, PUB41(ΔU)-eGFP is found at the anticlinal faces of the epidermal trichoblast cells (**Figure 1D-E**), a pattern resembling that of PIN proteins. In particular, PIN2 is localized in the root epidermis and affects root hair development and gravitropism (Rigas *et al*., 2013). However, the gravitropic response of the *pub41* mutants is similar to that of the WT plants (**Figure 1D-E**). This is in contrast to the *pin2* mutants, which show both affected root hair development and gravitropism (Rigas *et al*., 2013).

**Figure 1.**
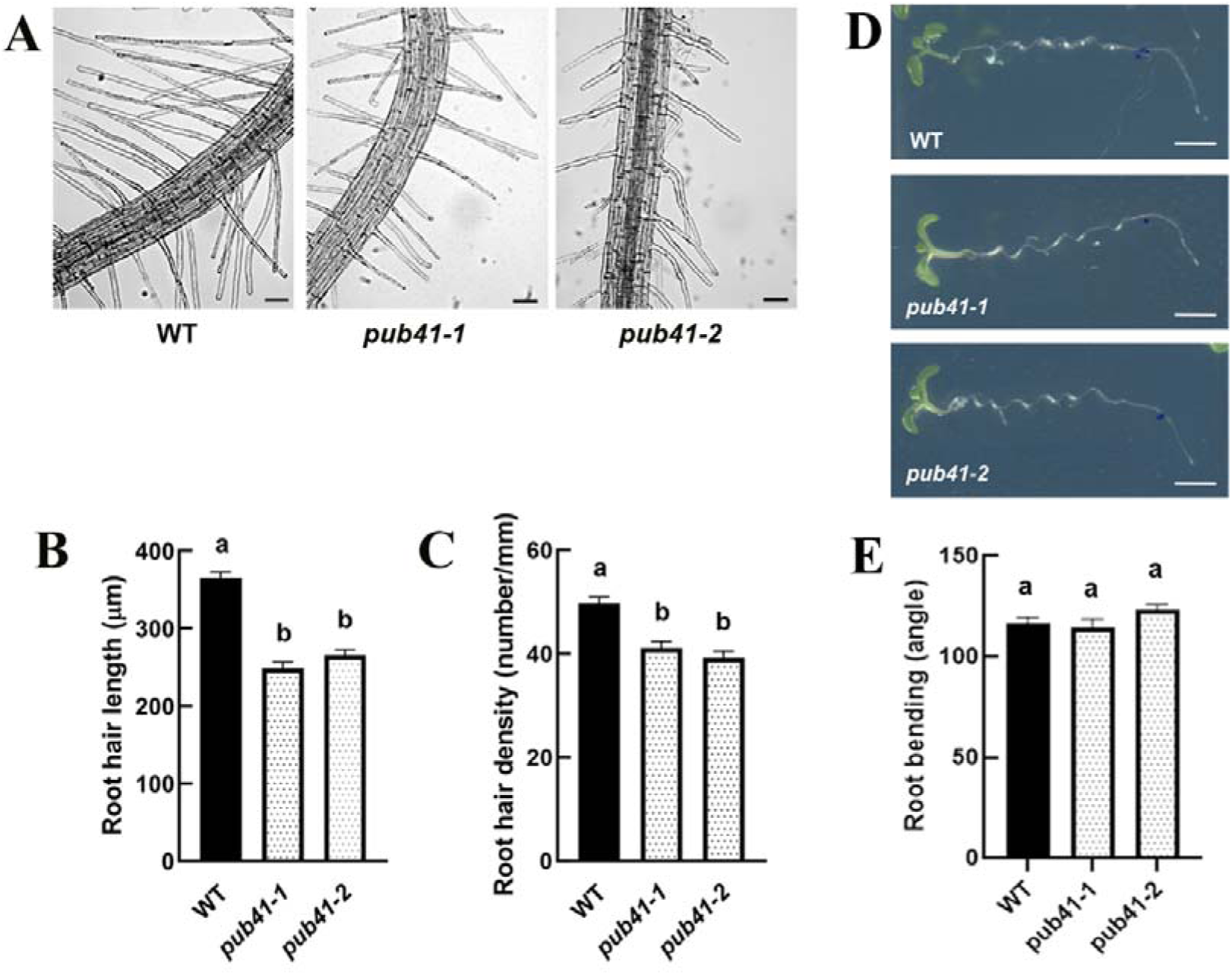
*pub41* mutants are affected in root-hair development but not in the gravitropic response. Ten-day-old Arabidopsis seedlings of the indicated genotypes were grown under standard conditions. (**A)** Representative images of the root differentiation zone of the primary root of ten-day-old seedlings. Scale bar = 50 μm. **(B-C).** The length (**B**) and density **(C)** of fully expanded root hairs in a 1-mm section of the root differentiation zone were determined. (**D**) Root gravitropism. Plates with 4-day old vertically grown seedlings were rotated by 90°. Seedlings were imaged 12 h later. Scale bar = 3 mm. (**E**) Quantitative analysis of root gravitropic response. Bars with different letters indicate significant differences (Tukey’s HSD post-hoc test p<0.01).

### Overexpressing *PUB41(ΔU)-eGFP* complements the root hair phenotype of *pub41* mutants

The U-box motif of E3 ligases binds the E2 enzymes (Trujillo, 2018). Thus, decoy E3s lacking the U-box and thus ubiquitin-conjugating activity can be used to study their protein targets, as the interaction with substrates does not result in the degradation of the target proteins (Lee *et al*., 2018). We constructed the pCAMBIA-derived *CaMV35S::PUB41(ΔU)-eGFP* plasmid, which encodes a truncated PUB41 protein lacking the U-box domain (residues 1-121), with the Met122 codon serving as the translation start of the truncated PUB41 protein. This plasmid was used to transform *pub41* mutant plants to circumvent background expression of the endogenous *PUB41* in the parental line. Homozygous transgenic plants, each resulting from a single T-DNA insertion event, were selected. Surprisingly, examination of the root hairs of three independent transgenic lines showed that the decoy *CaMV35S::PUB41(ΔU)-eGFP* complemented the root-hair phenotype of the *pub41-1* mutant, and enhanced root-hair length and density above that of the WT plants (**Figure 2A-C**). The high expression level of *PUB41(ΔU)-eGFP* transcripts was directly confirmed by RT-qPCR analysis in 5 tested lines, and revealed, as expected, that the steady state levels of the *PUB41(ΔU)-eGFP* transcripts driven by the highly active CaMV 35S promoter, were at least 10-fold higher than that of endogenous *PUB41* transcripts (**Figure 2D**). This major difference in expression levels accounts for *pub41* plants expressing *CaMV35S::PUB41(DU)-eGFP* developing more and longer root hairs than WT plants.

**Figure 2.**
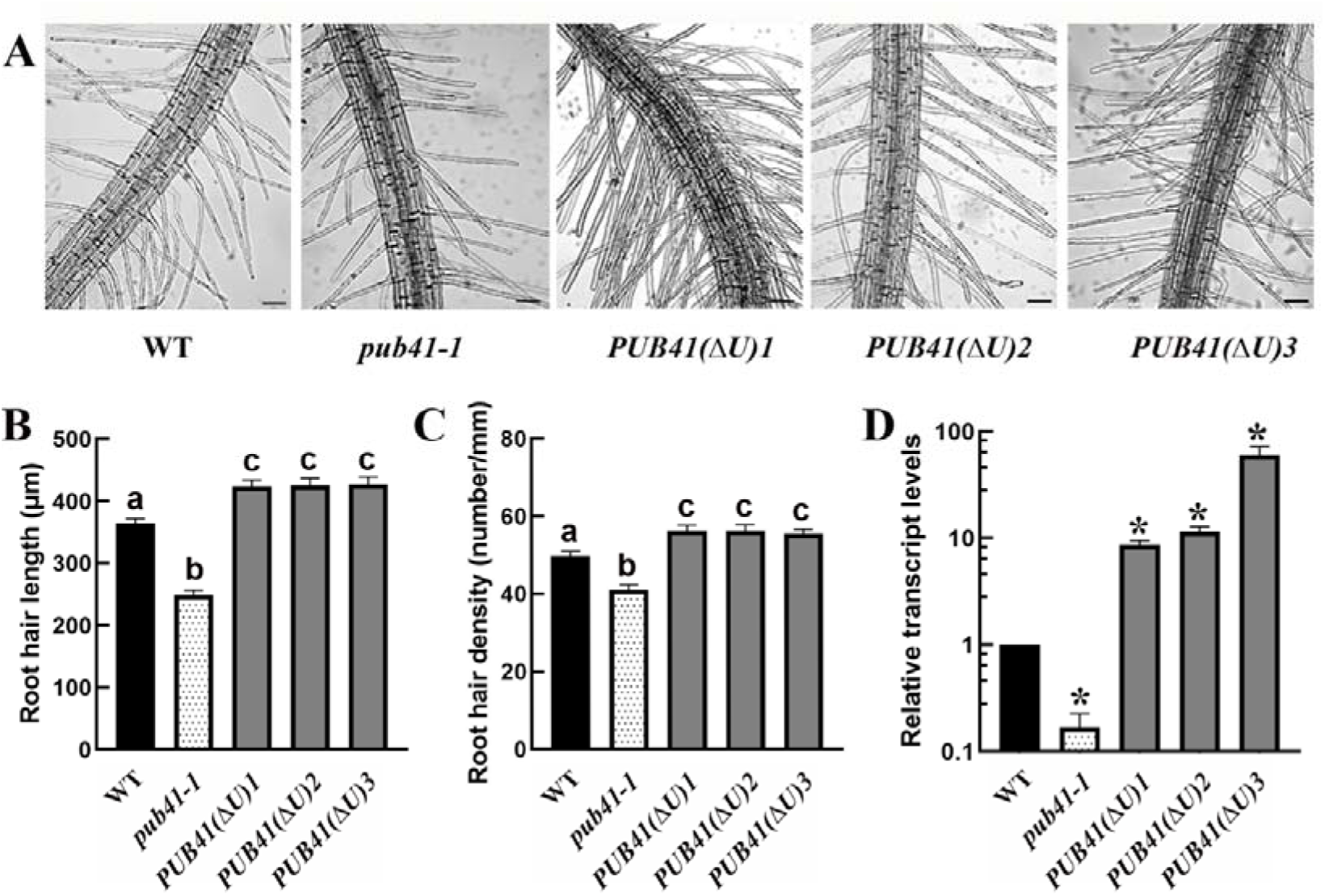
*PUB41*(ΔU) complements the *pub41-1* mutant root hair length and density phenotype. (**A)** Root hair phenotypes of WT, *pub41-1* mutant, and three independent lines of *pub41-1* expressing *35S::PUB41(ΔU)-eGFP*. Scale bar = 50 μm. **(B-C)** Quantitative analysis of root hair lengths and density. Bars with different letters indicate statistically significant differences (Tukey’s HSD post-hoc test p<0.01). (**D**) RT-qPCR analysis of *PUB41* expression. Asterisks denote the statistically significant differences between the mutant and overexpressing lines compared with WT plants (two-tailed paired Student’s t-test (p ≤ 0.01).

### Auxin modulates *PUB41* transcript and protein levels

Root hair development is modulated primarily by auxin, and we assayed whether exogenous auxin affects *PUB41* promoter activity and the levels of *PUB41* transcript and protein. Auxin-treated WT plants were assayed for the relative steady-state *PUB41* transcript levels in the roots by RT-qPCR. Indole-3-acetic acid (IAA) treatment increased *PUB41* transcript levels approximately two-fold (**Figure 3A**). To directly assess promoter activity, we expressed *PUB41::GUS* in seedlings and assayed GUS activity in roots of WT and IAA-treated plants. Treatment with 10 µM of IAA led to increased GUS activity, particularly in the root elongation and differentiation zones, indicating an auxin-dependent activation of the *PUB41* promoter (**Figure 3B**). In addition, treatment of *35S::PUB41(ΔU)-eGFP* plants with IAA resulted in elevated levels of the PUB41(ΔU)-eGFP) protein, predominantly in the root tip (**Figure 3C-D**). Thus, auxin positively regulates *PUB41* expression at both the RNA and protein levels.

**Figure 3.**
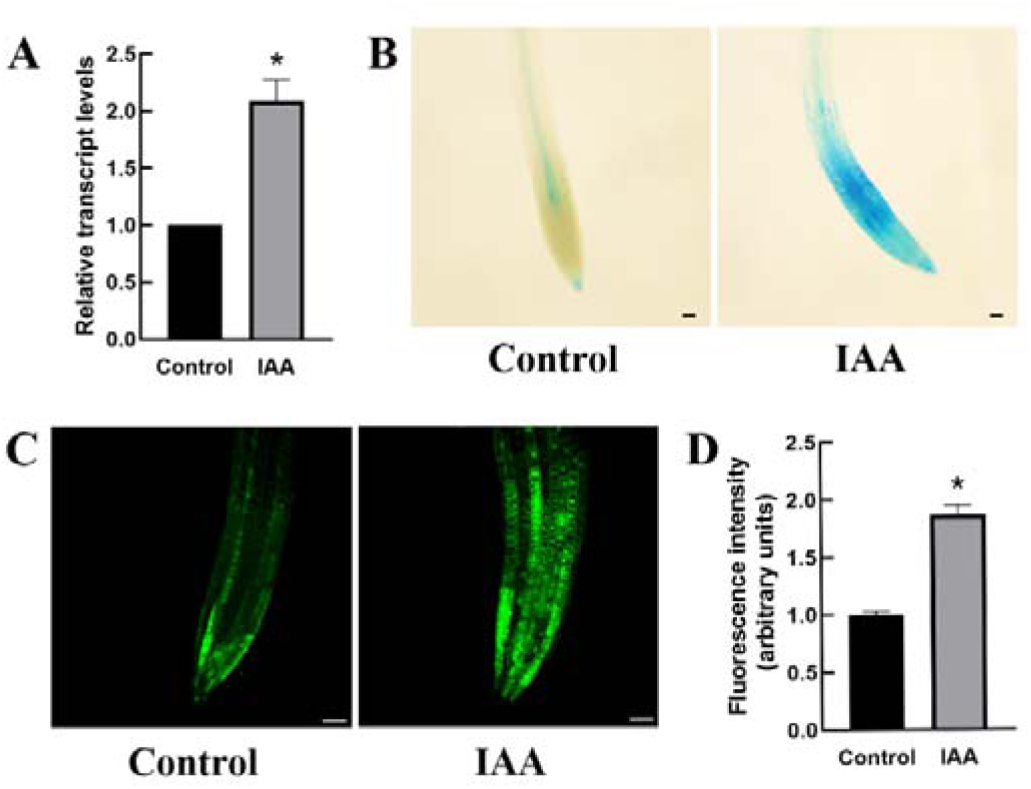
Auxin modulates *PUB41* RNA and protein levels. **(A)** RT-qPCR analysis of *PUB41* transcript levels in the roots of 10-day old WT seedlings untreated or treated with 10 µM IAA for 8 h. Data presented are the mean ± SE of 5 independent experiments. **(B)** Primary root tips of 8-day old WT seedlings expressing *PUB41::GUS,* untreated or treated with 10 µM IAA. Scale bar = 50 μm. **(C)** Primary root tips of transgenic plants expressing *35S::PUB41(ΔU)-eGFP* treated with 10 µM IAA. Scale bar = 50 µm). **(D)** Quantification of the GFP relative fluorescence intensity of 10-12 roots. In each experiment, IAA treatment was for 8 h and was repeated at least 3 times. Asterisks denote statistical significance as determined by a two-tailed paired Student’s t-test (p ≤ 0.01).

### Exogenous auxin induces root hair development in the elongation zone of WT, *pub41* mutants, and *PUB41(ΔU)-eGFP* expressing plants

To test whether overexpressing *PUB41* affects the auxin-sensing ability of the roots, we incubated seven-day-old WT, *pub41* mutants, and *pub41* mutants expressing *PUB41(ΔU)-*eGFP with 10 μM IAA. We examined the younger region near the root tip, that did not yet have root hairs in control plants of all genotypes. Although all genotypes developed root hairs 12-14 h following transfer to plates with 10 µM IAA (**Figure 4**), they differed in the extent of root hair number, with the lowest and highest numbers in *pub41-1* mutants, and *PUB41(ΔU)-eGFP* overexpressing plants, respectively.

**Figure 4.**
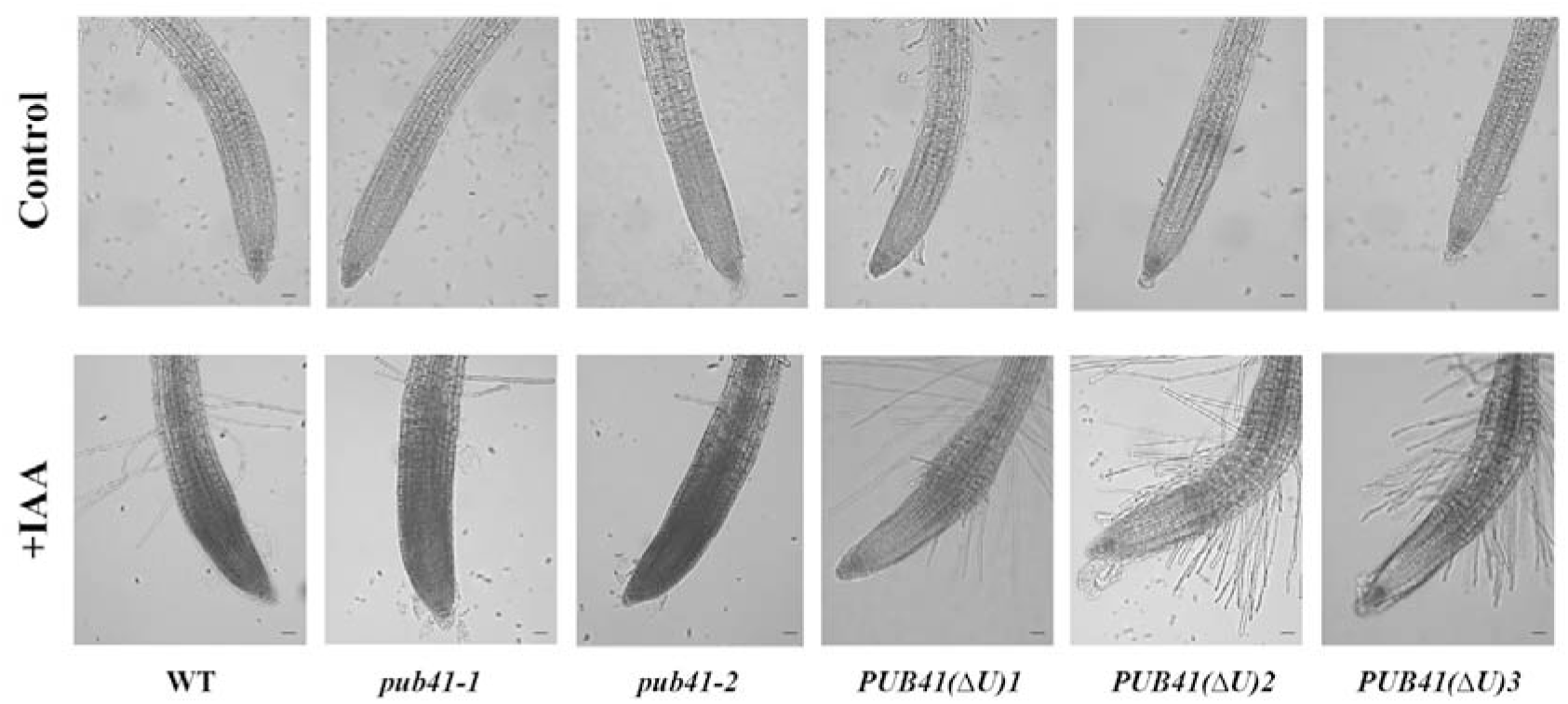
Induction of root hair formation by exogenous auxin. Seven-day-old seedlings of *pub41* mutants, WT and *PUB41(ΔU)-eGFP* overexpressing lines, were transferred to plates with growth medium lacking (control) or supplemented with 10 µM IAA (+IAA). Plates were incubated overnight under continuous light before the roots were examined. Scale bar = 50 μm.

### PUB41 protein localization is dependent on cell type and developmental stage

We observed PUB41(ΔU)-eGFP protein in all the root meristem cells, localized throughout the nuclei and cytoplasm (**Figure 5A-C**). However, in the elongation and differentiation zones, we found it mainly in the epidermis, primarily in trichoblasts and root hairs, but not in atricholblasts despite the CaMV 35S promoter being active in all cell types (**Figure 5D-E**). In all cell types, we observed PUB41(ΔU)-eGFP in both the nucleus and cytoplasm, however, within the cytoplasm its localization changed according to the developmental stage. In young newly divided nondifferentiated cells, PUB41(ΔU)-eGFP was evenly distributed in the cytoplasm (**Figure 5A**), similar to the pattern following transient expression of *PUB41-eGFP* and *PUB41(ΔU)-eGFP* in *N. benthamiana* leaves (Goldfarb, 2019; Sharma *et al*., 2024). In contrast, in epidermal cells of the elongation and differentiation zones, the protein was asymmetrically localized, preferentially at the anticlinal faces (perpendicular to the root axis) (**Figure 5D-E**). This indicates that the tissue- and subcellular localization of PUB41 are post-translationally regulated in a cell-type-specific manner.

**Figure 5.**
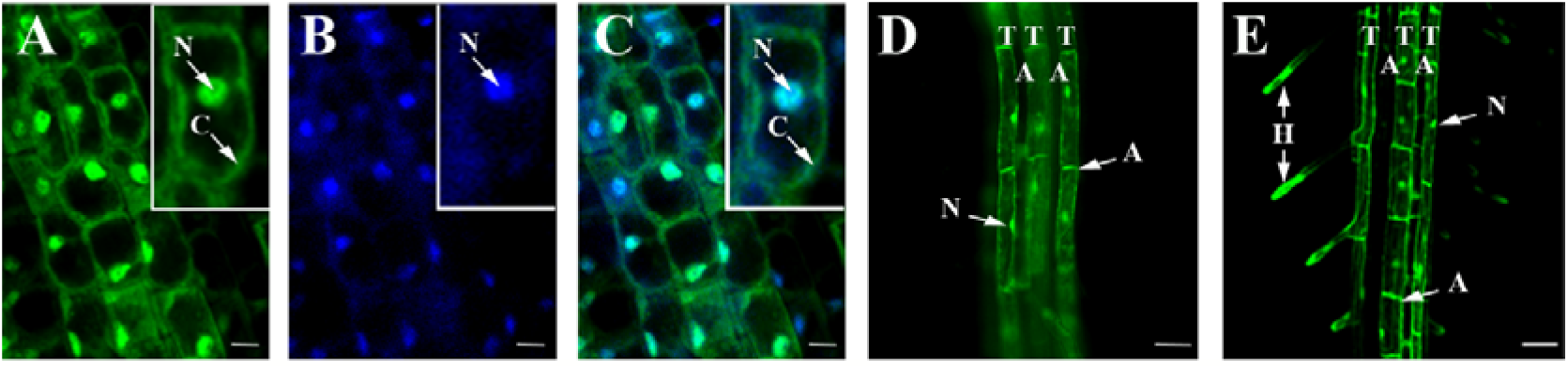
Subcellular localization of PUB41(ΔU)-eGFP in the roots. (**A-C**) Roots of one-week-old seedlings were counterstained with the DNA marker DAPI and meristematic zones were analyzed for eGFP (**A**) and DAPI (**B**) fluorescence. (**C**) merged image of (**A**) and (**B**), arrows marked N and C indicate nuclei and cytoplasm, respectively. Scale bars = 10 μm. (**D-E**) Elongation and differentiation zones, respectively, of two-week-old plants showing eGFP fluorescence. Arrows indicate: H, root hairs; T, trichoblasts; A, atrichoblasts; N, nuclei; A, anticlinal face of the planar cell. Scale bar = 50 μm.

### PUB41(ΔU) interacts *in vivo* and *in vitro* with PIN2

The localization pattern of PUB41(ΔU)-eGFP at the anticlinal faces of root trichoblasts resembles that of the auxin transporter PIN2, which is required for root hair development (Rigas *et al*., 2013). To test if PUB41 interacts with PIN2 *in planta*, we crossed homozygous *35S::PUB41(ΔU)-eGFP* Arabidopsis plants with homozygous *PIN2::PIN2-mCherry* plants. Confocal microscopy analysis of roots of the hybrid plants confirmed the colocalization of these proteins, validating the interaction between PUB41 and PIN2 (**Figure 6A**).

**Figure 6.**
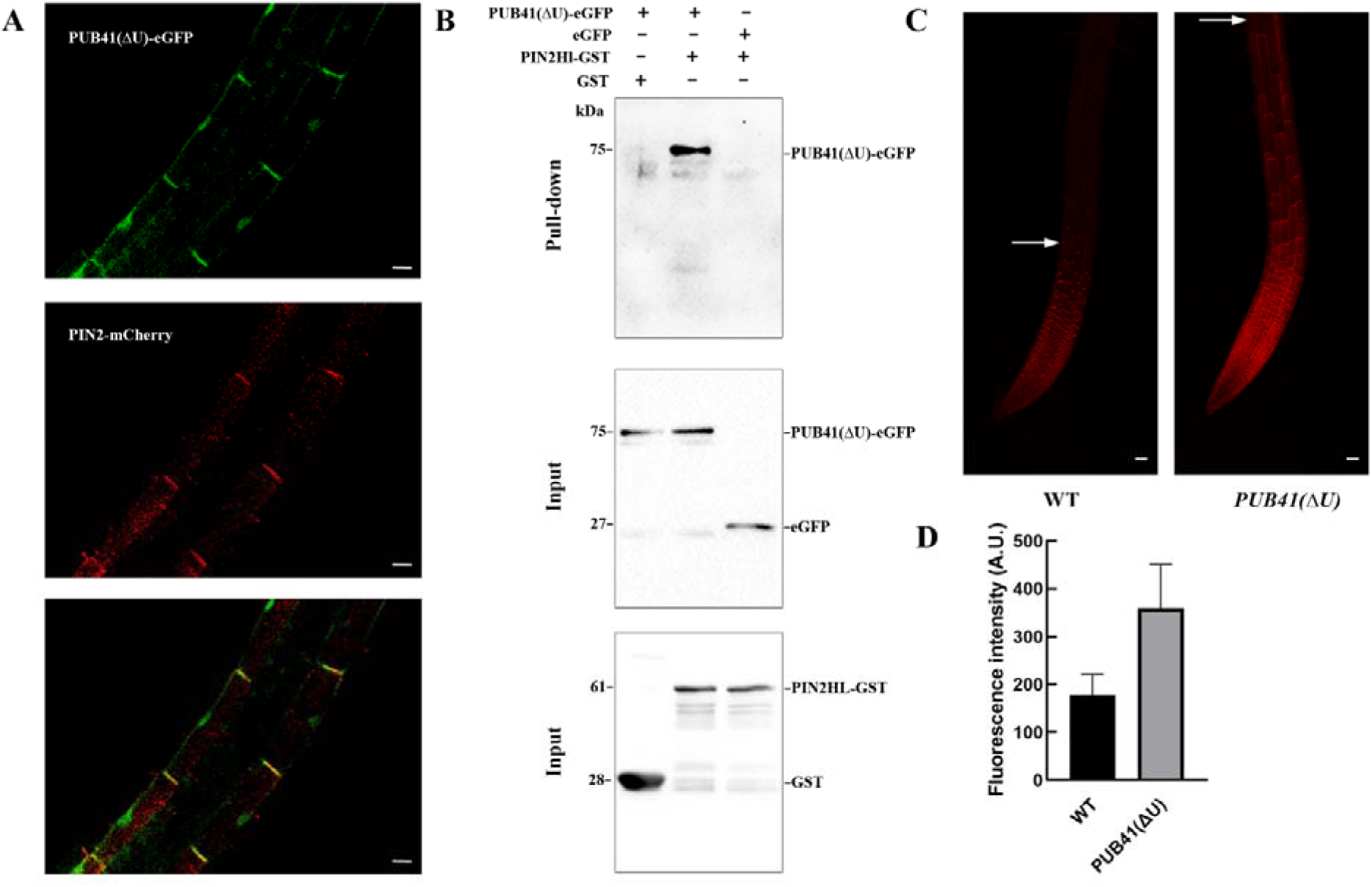
PUB41 interacts with PIN2. **(A)** Co-localization of PUB41(ΔU)-GFP and PIN2-mCherry in epidermal cells of primary root elongation zone. Scale bar = 10 μm. **(B)** Pull-down assay showing direct interaction between PUB41-eGFP and GST-PIN2HL. **(C)** PUB41 stabilizes the expression of PIN2 in the upper root elongation zone. Confocal microscope images showing the expression pattern of PIN2-mCherry protein in WT (top) and in *35S::PUB41(ΔU)-GFP* expressing plants (bottom). Scale bar = 50 µm. The experiment was repeated at least 3 times with 10 plants per genotype.

PUB41 is a water-soluble protein, whereas PIN2 is embedded in the cell membrane (above). To assay whether PUB41(ΔU) binds the PIN2 hydrophilic loop (PIN2HL), soluble protein fractions, extracted from Arabidopsis plants expressing PUB41(ΔU)-eGFP were incubated with *E. coli*-expressed GST-tagged PIN2HL in an *in-vitro* pull-down assay. Our result confirmed that PUB41(ΔU) binds PIN2HL (**Figure 6B**). This is compatible with the *in planta* colocalization of the proteins (**Figure 6A**).

Furthermore, microscopic analysis of plants expressing PIN2-mCherry in the background of PUB41(ΔU)-GFP revealed an extended expression pattern of PIN2-mCherry in the root elongation zone compared with plants expressing PIN2-mCherry in a WT background (**Figure 6C**). This suggests that decoy PUB41(ΔU) binding of the PIN2HL may contribute to stabilizing the auxin transporter.

### Full-length PUB41 interacts with PIN2 and complements the *pub41* mutant

Transient expression of PUB41-eGFP in Arabidopsis seedlings expressing PIN2-mCherry, showed that these proteins are colocalized *in planta* at the horizontal faces of the root cells (**Figure 7A-C**). Overexpressing full-length *PUB41* complemented the root hair development phenotype of the *pub41-1* mutant (**Figure 7D-G**), as did overexpressing the decoy *PUB41(ΔU)-eGFP* (**Figure 2**).

**Figure 7.**
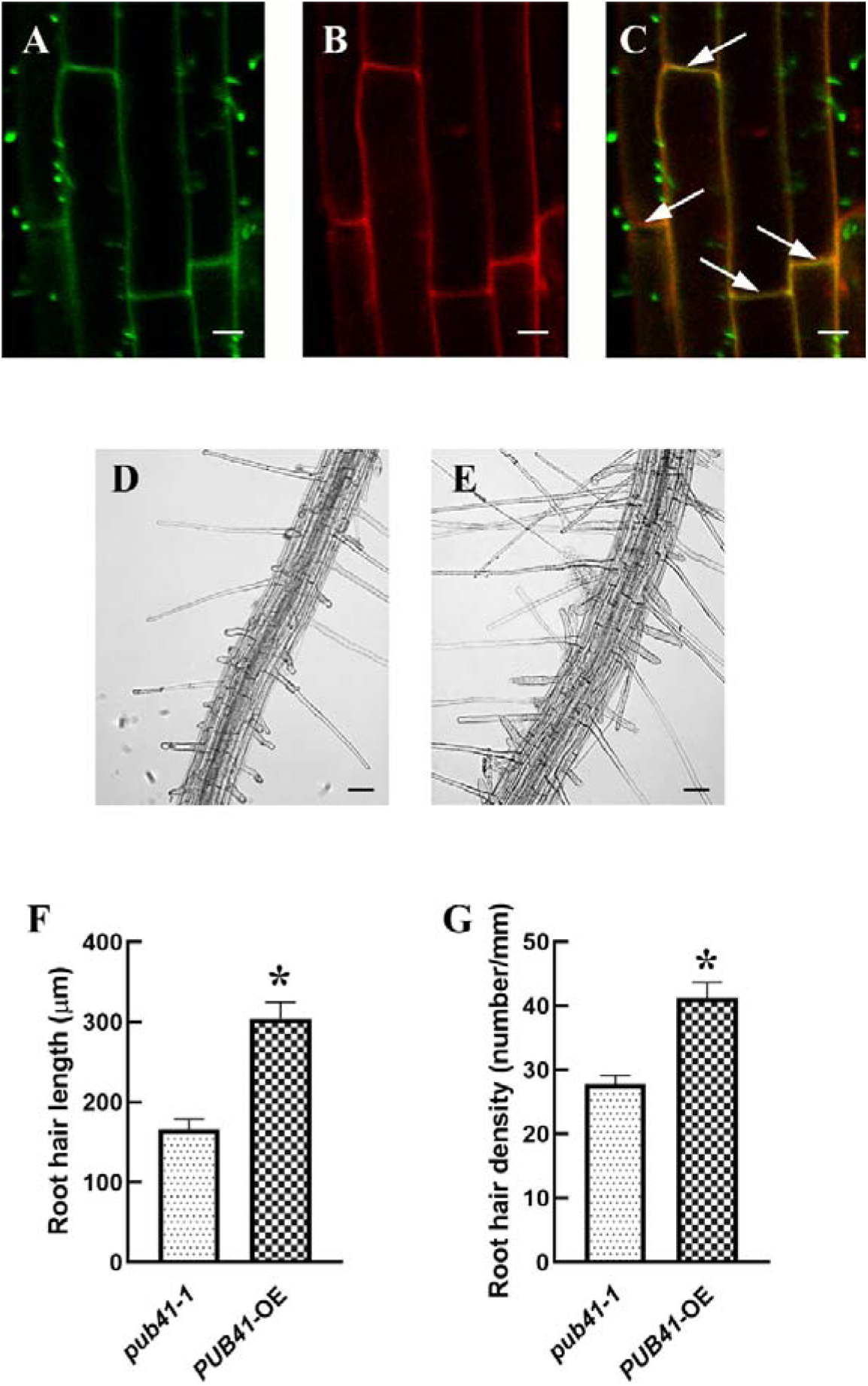
Full-length PUB41 interacts with PIN2 and complements the *pub41* root hair phenotype. **(A-C)** Roots of *PIN2::PIN2-mCherry* Arabidopsis seedlings transiently expressing *CaMV 35S::PUB41-eGFP.* Roots were examined by confocal microscope: (**A**) PUB41-eGFP; (**B**) PIN2-mCherry; (**C**) Merged. Scale bars = 10 μm. (**D-G**) Root hair phenotype of the *pub41-1* mutant (**D**), and *pub41-1* mutant plants transformed with *CaMV 35S::PUB41-eGFP* (**E**). Scale bar = 50 μm. **(F, G)** Quantitative analysis of root hair lengths and density. Each experiment was carried out 3 times with at least 10 plants for each genotype. Asterisks denote the statistically significant differences between mutant and overexpressing lines compared with WT plants (two-tailed paired Student’s t-test (p ≤ 0.01).

## Discussion

We recently showed that *PUB41* is essential for drought resilience and seed dormancy in Arabidopsis and that *PUB41* transcript levels are significantly higher in roots than in shoots, implying a root-specific function for this gene. Indeed, we observed high *PUB41* promoter activity in trichoblasts and root hair cells suggesting that PUB41 accumulation in these cells is regulated at the RNA transcript and protein levels and that PUB41 may be involved in root hair development (Sharma *et al*., 2024).

Here we show that indeed *pub41* mutants developed fewer and shorter root hairs compared with WT plants and that *PUB41* promoter activity is increased particularly in mature root hairs of the differentiation zone and the collet region (**Figure 1**). Treatment with IAA leads to a doubling of the steady-state levels of *PUB41* transcripts in roots, compatible with increased promoter activity and compatible with a role in root hair development (**Figure** 3**A-B**). Furthermore, the PUB41 protein was stabilized by auxin treatment (**Figure** 3**C-D**).

Here, using inactive decoy PUB41(ΔU), we generated transgenic Arabidopsis plants overexpressing PUB41(ΔU)-eGFP to study its localization and to identify interacting proteins. In the root meristem, PUB41(ΔU)-eGFP is evenly distributed in both the nucleus and the cytoplasm (**Figure 5**), similar to PUB41-eGFP transiently expressed in leaf epidermal cells *of N. benthamiana* (Sharma *et al*., 2024). However, in the elongation and differentiation zones, the protein, while present in both nucleus and cytoplasm, becomes asymmetrically localized in the cytoplasm and concentrated at the anticlinal faces of the cells (**Figure 5**). Subcellular localization is primarily determined by a signal peptide/localization sequence but can be modulated by post-translational modifications (PTM) such as phosphorylation, acetylation, SUMOylation, and ubiquitylation (Zhang and Zeng, 2020). Moreover, crosstalk between ubiquitylation and such PTMs for example, phosphorylation, may alter the recognition site of the E3 ligase and affect protein stability (Lee *et al*., 2023). In the differentiation zone of the root, we detected PUB41(ΔU)-eGFP protein only in trichoblasts and root hairs, but not in atrichoblasts (**Figure 5D-E**). Our interpretation is that the degradation of PUB41 is highly regulated and cell-type dependent, with PUB41 being more accessible to degradation in atrichoblasts than in trichoblasts. In the differentiation root region, we observed PUB41(ΔU)-eGFP at the apical faces of the epidermal cells (**Figure 5D-E**). This subcellular distribution coincides with that of the PIN auxin transporter proteins. The similarity between the root hair phenotype of *pub41* and that of *pin2* mutants (Rigas *et al*., 2013) suggests that these two proteins may interact. Indeed, we found that PUB41 colocalized with PIN2 in the differentiation zone of the roots (**Figure 6A**) and showed by *in vitro* co-precipitation that PUB41 binds the cytoplasmic hydrophilic loop of PIN2 (**Figure 6B**). This colocalization may result from direct assignment of PUB41 to the apical faces of epidermal cells, or from its protein-protein interaction with prelocalized PIN2. Binding of the decoy PUB41(ΔU)-eGFP to PIN2 resulted in enhanced PIN2 levels and extended the root region in which it is found (**Figure 6C**). These enhanced PIN2 levels are predicted to result in increase of local auxin levels that would enhance the emergence and growth of root hairs. Whereas *pin2* mutant plants show impaired root hair development and gravity response (Rigas *et al*., 2013), *pub41* mutant plants show only impaired root hair development, but their gravitropism is unaffected (**Figure 1**). This suggests that PIN2 affects root hair development and gravitropism via two different pathways. In *pub41* mutants overexpressing either catalytically active *PUB41* or the inactive decoy *PUB41(ΔU)* restored root hair development (**Figures 4-7)**. We propose that the PUB41 Armadillo motifs interact with the PIN2 hydrophilic loop and stabilize the auxin transporter. One possibility is that PUB41 binds an amino acid sequence within the cytoplasmic loop of PIN2 that also acts as the recognition site for a different E3 ligase involved in PIN2 degradation (Pan *et al*., 2009; Konstantinova *et al*., 2022; Retzer *et al*., 2022). The hydrophilic loop of canonical PINs, including PIN2, possesses many residues that can be phosphorylated and ubiquitylated (reviewed by Zwiewka *et al*., 2019). These PTMs affect the subcellular localization of PINs, their stability, and probably their interactions with other proteins (Zwiewka *et al*., 2019). PIN2 is localized at the apical faces of root epidermis and the basal faces of root cortex cells, a distribution modulated by the phosphorylation status of the hydrophilic loop (Michniewicz *et al*., 2007). Whereas protein ubiquitylation most commonly results in destabilizing the modified protein, emerging evidence suggests that E3 ligases can also contribute to protein stabilization (Abu Ahmad *et al*., 2021; Hong *et al*., 2022). Our results resemble those described for SAV4 (Chen *et al*., 2022), where SAV4 colocalizes with PIN2, and interacts with its hydrophilic loop. SAV4 enhances PIN2 membrane clustering and stability through direct protein-protein interaction. Interestingly, SAV4 possesses a single ARM motif (residues 120-153), compared with the 5 ARM motifs in PUB41. Moreover, auxin enhanced activity of the plasma-membrane-associated TMK1 kinase phosphorylates and stabilizes PIN2 (Rodriguez *et al*., 2025).

PIN2 also modulates the root gravitropic response (Rigas *et al*., 2013). Surprisingly, elimination of *PUB41* did not affect the root gravitropic response (**Figure 1D-E**), suggesting that the effects of PIN2 on root hair development and gravitropic response can be uncoupled. This explanation is supported by (Templalexis *et al*., 2021): Mutating the Arabidopsis TINY ROOT HAIR (TRH1) gene, encoding K+ transporter, impairs root hair development and the root gravitropic response (Rigas *et al*., 2001; Vicente-Agullo *et al*., 2004). However, using an elegant approach expressing the TRH1 gene driven by root cell type-specific promoters, Templalexis et al uncoupled the auxin-mediated root-hair development and gravitropism activities. Both the root-hair and gravitropic response phenotypes of thr1 could be complemented by driving expression of THR1 by its endogenous promoter. In contrast, either the root-hair development or the gravitropic response could be complemented by driving the expression of the *TRH1* gene with the *PIN2* or *PIN1* promoters, respectively (Templalexis *et al*., 2021). These results suggest the epidermal expression of *TRH1* is essential for root hair development, whereas expression in the central cylinder is required for its effect on gravitropic response. In the differentiation zone, PUB41(ΔU)-eGFP is primarily present in the root epidermis (**Figure 5D-E**), thus affecting root hair development but not the gravitropic response.

The detailed mechanism controlling PUB41 stabilization by auxin, and the interaction of PUB41 with PIN2 and the other E3s, remains to be elucidated.

## Materials and methods

### Plant Material

The T-DNA insertion lines *pub41-1* (SALK_099012) and *pub41-2* (SALK_142012) (Sharma *et al*., 2024) were acquired from the Arabidopsis Resource Center, Columbus, Ohio. The seeds of Arabidopsis wild-type (Columbia ecotype) plants expressing the *PIN2:PIN2-mCherry* construct (Zhang *et al*., 2020) were generously gifted by Prof. Jiří Friml (Institute of Science and Technology, Austria). Arabidopsis plants expressing *35S::PUB41-eGFP* and *PUB41::GUS* were described earlier (Sharma *et al*., 2024). The 35S::PUB41(Δ U-box)-eGFP plasmid was constructed by amplifying the DNA sequence encoding amino acid residues Met122-Phe559 using primers containing linkers with the *Nco*I and *Pst*I restriction sites (Table S1). This sequence was subcloned into the respective restriction sites of the pGA-eGFP3 vector (Maymon *et al*., 2022) in frame with the sequence encoding eGFP. The resulting pCAMBIA plasmids were introduced into *Agrobacterium tumefaciens* GV3101, which was used to transform *pub41-1* Arabidopsis mutant plants by the floral dip method (Clough and Bent, 1998). Transgenic plants were selected on hygromycin-containing Arabidopsis growth medium.

Plants co-expressing *PIN2::PIN2mCherry* and *35S::PUB41(*Δ*U)-eGFP* were generated by crossing homozygous plants expressing the individual constructs, where pollen collected from *35S::PUB41(*Δ*U)-eGFP* plants was used to fertilize plants expressing *PIN2::PIN2mCherry*. The seeds were collected and plated on hygromycin-containing growth medium to select plants expressing both constructs. Roots co-expressing *PIN2::PIN2mCherry* and *35S::PUB41-GFP* were also obtained by transient expression of *35S::PUB41-GFP* in roots of *PIN2::PIN2mCherry* plants. Cultures of *Agrobacterium* GV3101 harboring pCAMBIA *35S::PUB41-eGFP* or p19 plasmids were grown to OD_600_ =0.6. Bacteria were collected by centrifugation and resuspended in resuspension buffer (0.25 x MS, 100 µM acetosyringone, 0.005% Silwet) to yield OD_600_ =1. One ml of each bacterial suspension was added to 2 ml of resuspension buffer and incubated overnight. A 2 ml aliquot of the bacterial mix was added to a small Petri dish containing seven-day-old *PIN2::PIN2-*mCherry seedlings. Plants were incubated in the dark for 12 h, followed by several washes using the resuspension buffer without Silwet. Roots were harvested and examined by confocal fluorescent microscopy.

### Plant growth

Arabidopsis plants were cultivated at 23°C, under 50% relative humidity, and a photoperiod of 12 h light and 12 h darkness. Seeds were surface sterilized and imbibed at 4 °C for at least four days before being sown either on a solid medium [0.5× Murashige and Skoog (MS), 0.5% plant agar, and 0.5% (w/v) sucrose], or in pots as described previously (Sharma *et al*., 2024). Auxin (IAA) was applied by incubating ten-day old plate-grown seedlings on Whatman No. 1 filter paper soaked in 0.5 x MS and 0.5% (w/v) sucrose with the indicated hormone concentration.

### Root hair length and density

Surface-sterilized stratified seeds of the indicated genotypes were plated on agar-solidified growth medium. Plates were incubated vertically, and plants were harvested 10 days later. For phenotyping root hair traits in plants expressing *35S::PUB41-eGFP*, seeds of the T0 generation were plated on agar-solidified medium supplemented with 30 μg/ml hygromycin and incubated in the dark for 3 days. Seeds of the parental *pub41-1* mutant line were plated and grown in parallel on medium lacking antibiotics. The germinated etiolated seedlings were then transferred to fresh Petri dishes containing antibiotic-free medium, and the plates were kept 5 more days in a vertical position. Seedlings were harvested, and root differentiation zones were imaged using a CCD camera (DS-Fi1; Nikon) mounted on the eyepiece tube of Nikon Eclipse Ci microscope. For root hair density, the number of root hair cells was counted in a 1 mm section of the differentiation zone where trichoblasts had fully expanded (Sánchez-Fernández *et al*., 1997). Root hair number and length were measured in at least ten seedlings per biological replica (Sánchez-Fernández *et al*., 1997) using Nikon NIS elements in-built measurement tools.

### Root gravitropism

Surface-sterilized stratified seeds of the indicated genotypes were plated on agar-solidified growth medium. Plates were kept vertically for 4 days and then turned by 90°. Plates were then imaged using an Epson Perfection V370 photo Scanner, and root bending angles were determined using FIJI (Schindelin *et al*., 2012)

### GUS staining

GUS staining was performed as described (Sharma *et al*., 2024). Each experiment was repeated three times using three independent homozygous lines.

### RT-qPCR analysis

Arabidopsis plants of indicated genotypes and age were plated on top of a silk print mesh (Eisner *et al*., 2021) in plates containing agar-solidified medium supplemented with 0.5× Murashige and Skoog (MS) and 0.5% (w/v) sucrose. Mesh-grown plants were transferred to plates containing Whatman No. 1 paper soaked in 0.5 x MS+0.5% (w/v) sucrose, with or without the indicated reagent concentration and incubated for the specified time, before harvesting the roots. RNA isolation, cDNA synthesis, primer design, and RT-qPCR assays for determining relative steady-state transcript levels were performed as described (Maymon *et al*., 2022). Primers used in this study are listed in Supplementary Table S1.

### Recombinant protein expression in *E. coli*

The DNA sequence encoding the cytoplasmic hydrophilic loop of PIN2 (PIN2-HL, residues 153-503, (Chen *et al*., 2022)) was amplified from WT plant cDNA using a primer set containing *NcoI* and *XbaI* linkers (Table S1). The resulting DNA fragment was cloned into the respective sites of the pGST-parallel2 vector (Sheffield *et al*., 1999) using the restriction-free (RF) cloning method (Bond and Naus, 2012). The pGST-Parrallel2 vector and the recombinant pGST-PIN2-HL-parralel2 plasmid were introduced into *E. coli* BL21(DE3)pLysS cells. Mid-log phase cultures (OD_600_= 0.6) were induced at 16 °C by adding 0.6 mM IPTG.

### Pull-down assay

IPTG-induced bacterial cultures were centrifuged, and the harvested cells resuspended in solubilization buffer (20 mM Tris, 200 mM NaCl, (pH 7.6), 1X protease inhibitor cocktail (BiMake)) supplemented with 1 mg/ml lysozyme, using a 1 gr pellet/ 5 ml buffer ratio. The bacterial suspension was passed a few times through a syringe equipped with a 0.8 x 38 mm needle, followed by sonication. The lysate was then centrifuged at 12,000 x g rpm for 15 min at 4 °C, and supernatants containing recombinant GST-PIN2HL and GST-tag were incubated at 4 °C for 1.5 h under continuous agitation. Bead-bound recombinant proteins were washed several times with solubilization buffer lacking lysozyme, and protein-bound resins were aliquoted (30 µl each) and stored at -80 °C for later *in vitro* pull-down assays.

Arabidopsis seeds of the indicated lines expressing *PUB41(*Δ*U)-eGFP*, *eGFP tag*, and WT, were plated on agar-solidified growth medium. The Petri dishes were covered with aluminum foil and incubated in the growth room for 6 days. Seedlings were harvested and homogenized in liquid nitrogen using a mortar and pestle. Homogenization buffer [20 mM Tris, (pH 7.6), 200 mM NaCl, protease inhibitor cocktail (BiMake), and 0.2% NP40] was added at a 10 ml buffer / g seedlings ratio. Homogenates were centrifuged at 12,000 rpm, 4 °C, for 15 min, and supernatants were collected. Before the pull-down assay, total protein concentrations were estimated by both nanodrop (NanoDrop ND-1000; NanoDrop Technologies), and GFP levels were assayed by western blotting.

For the assay, 0.7 ml aliquots of plant extracts were added to tubes containing 30 μl of protein-loaded glutathione resin and incubated for 90 min at 4 °C with continuous agitation. Beads were collected by centrifugation for 1 min, 600 x g at 4 °C, supernatants were discarded, and beads were washed 3 times in the same buffer. Beads were resuspended in Laemmli’s SDS PAGE sample buffer (Laemmli, 1970), and incubated in a boiling water bath for 8 minutes. Proteins were resolved by SDS-PAGE, and electroblotted onto nitrocellulose membranes. Membranes were stained with Ponceau S, and recombinant proteins were detected using monoclonal primary antibodies: anti-GFP (ab1218, Abcam), anti-GST (G018, Applied Biological Materials Inc), and anti-His tag (G020, Applied Biological Materials Inc). Peroxidase-coupled anti-mouse IgG antibody (Sera Care 5450-0011) was used as a secondary antibody. Membranes were then incubated in a reaction mix prepared using luminol and peroxide solutions provided in the WesternBright ECL (Advansta) kit, and chemiluminescent signals were recorded with ImageQuant RT ECL Imager (GE Healthcare).

### Microscopy

Light microscopy was performed using a Nikon SMZ100 stereoscope. Fluorescent images were taken with a Nikon Eclipse Ci-L fluorescence microscope or Zeiss LSM-880 (Axio Observer Z1, inverted) laser scanning microscope equipped with Definite Focus 2 and the Airyscan detector. The following excitation and emission wavelengths were used: for GFP (Ex/Em = 488/507 nm), for mCherry Ex/Em =587/610 nm), for DAPI (Ex/Em = 405/470 nm). Images were processed using Fiji/ImageJ (Schindelin et al., 2012). For each experiment, all images were captured with identical microscope equipment, camera configuration, and exposure durations.

### Statistical analysis

All experiments were carried out with at least 3 biological repeats. Statistical analyses were conducted using PRISM version 10.0.0 (GrapPad Software, Boston, Massachusetts USA) on Windows 10 (post-hoc Tukey’s HSD test, Kruskal-Wallis non-parametric test, student’s t-test). Data Differences were considered significant at P < 0.05.

## Acknowledgments

AS was partially supported by a fellowship from the Kreitman School of Advanced Research Studies, Ben-Gurion University of the Negev.

## Author Contributions

AS, DR, and DBZ designed the experiments, generated and analyzed the data, and wrote the manuscript. AS carried out the experiments. All authors read and approved the manuscript.

## Data availability statement

The data that support the findings of this study are available from the corresponding author upon reasonable request.

## Supporting Information

Additional Supporting Information may be found in the online version of this article.

**Table S1.** Primers used in this study.

