## Supplemental Table S1 for "Ubiquitin ligase PUB41 modulates root hair development in Arabidopsis via interaction with the auxin polar transporter PIN2"

**Supplementary Table S1. Primers used in this study.**

| Application | Sequence | Primer | No. |
| --- | --- | --- | --- |
| Constructing 35S::PUB41(ΔU)-eGFP | ACTACCATGGATAAAGATCCTAATCCAAGTC | PUB41(Δ Ubox) eGFP For | 1 |
| Constructing 35S::PUB41(ΔU)-eGFP | AACTGCAGAACTGGGAGGAATAAGCAAAAC | PUB41 eGFP Rev | 2 |
| Cloning of pGST-par2-PIN2HL | ACCTGTATTTTCAGGGCGCCATGGAGTTCCGTGGGGCTAAG | pGST_par2-PIN2_HL FOR | 3 |
| Cloning of pGST-par2-PIN2HL | GCTCGAGACTGCAGGCTCTAGAGTTAGGGTTTCGAATGAGTTTTCT | pGST_par2-PIN2_HL REV | 4 |
| qPCR in WT | GAGCCGCTTCTTCTTTCTTCATC | PUB41-FP2 | 5 |
| qPCR in WT | CGTTGGTGGAAAGAGAACCATC | PUB41-RP2 | 6 |
| qPCR of PUB41(ΔU) | TGTGTGGCGGTTCTGTTGAC | 41RT-FP-S2 | 7 |
| qPCR of PUB41(ΔU) | CTTCGACGCCTTCTCCTTCA | 41RT-FP-S2 | 8 |
| qPCR | CGACTTCTTCAAGTCCGCCA | eGFP-FP | 9 |
| qPCR | TCTTGTAGTTGCCGTCGTCC | eGFP-RP | 10 |
| qPCR | TCCCTCAGCACATTCCAGCAGAT | ACTIN2-FP | 11 |
| qPCR | AACGATTCCTGGACCTGCCTCATC | ACTIN2-RP | 12 |
